# Species-level microbiome composition of activated sludge - introducing the MiDAS 3 ecosystem-specific reference database and taxonomy

**DOI:** 10.1101/842393

**Authors:** Marta Nierychlo, Kasper Skytte Andersen, Yijuan Xu, Nick Green, Mads Albertsen, Morten S. Dueholm, Per Halkjær Nielsen

## Abstract

The function of microbial communities in wastewater treatment systems and anaerobic digesters is dictated by the physiological activity of its members and complex interactions between them. Since functional traits are often conserved at low taxonomic ranks (genus, species, strain), the development of high taxonomic resolution and reliable classification is the first crucial step towards understanding the role of microbes in any ecosystem. Here we present MiDAS 3, a comprehensive 16S rRNA gene reference database based on high-quality full-length sequences derived from activated sludge and anaerobic digester systems. The MiDAS 3 taxonomy proposes unique provisional names for all microorganisms down to species level. MiDAS 3 was applied for the detailed analysis of microbial communities in 20 Danish wastewater treatment plants with nutrient removal, sampled over 12 years, demonstrating community stability and many abundant core taxa. The top 50 most abundant species belonged to genera, of which >50% have no known function in the system, emphasizing the need for more efforts towards elucidating the role of important members of wastewater treatment ecosystems. The MiDAS 3 taxonomic database guided an update of the MiDAS Field Guide – an online resource linking the identity of microorganisms in wastewater treatment systems to available data related to their functional importance. The new field guide contains a complete list of genera (>1,800) and species (>4,200) found in activated sludge and anaerobic digesters. The identity of the microbes is linked to functional information, where available. The website also provides the possibility to BLAST the sequences against MiDAS 3 taxonomy directly online. The MiDAS Field Guide is a collaborative platform acting as an online knowledge repository and facilitating understanding of wastewater treatment ecosystem function.

## Introduction

Biological nutrient removal (BNR) plants are the principal wastewater treatment systems across the urbanized world. Besides their primary role in removing pollutants and pathogens, these systems are increasingly seen as resource recovery facilities, contributing to sustainable resource management. At-plant anaerobic digestion exploits the value of sludge as a source of energy, reducing plant carbon footprint and placing wastewater treatment plants (WWTPs) on the road to achieving a circular economy. Complex microbial communities define the function of biological wastewater treatment systems, and understanding the role of individual microbes requires knowledge of their identity, to which known and putative functions can be assigned. The gold standard for identification and characterization of the diversity, composition, and dynamics of microbial communities is presently 16S rRNA gene amplicon sequencing (Boughner and Singh, 2016). A crucial step is a reliable taxonomic classification, but this is strongly hampered by the lack of sufficient reference sequences in public databases for many important microbes present in wastewater treatment systems (McIlroy et al., 2015). In addition, high taxonomic resolution is missing from the large-scale public reference databases (SILVA (Quast et al., 2013), Greengenes (DeSantis et al., 2006), RDP (Cole et al., 2014)), often resulting in poor classification.

MiDAS (Microbial Database for Activated Sludge) was established in 2015 as an ecosystem-specific database for wastewater treatment systems that provided manually curated taxonomic assignment (MiDAS taxonomy 1.0) based on the SILVA database and associated physiological information profiles (midasfieldguide.org) for all abundant and process-critical genera in activated sludge (AS) (McIlroy et al., 2015). Both taxonomy and database were later updated (MiDAS 2.0) to cover abundant microorganisms found in anaerobic digesters (AD) and influent wastewater (McIlroy et al., 2017). However, more high-identity 16S rRNA gene reference sequences are needed to be able to classify microbes at the lowest possible taxonomic levels. Currently, taxonomic classification can only be provided down to genus level, limiting elucidation of the true diversity in these ecosystems. Consequently, physiological differences between the members of the same genus that coexist in activated sludge cannot be reliably linked to the individual phylotypes due to insufficient phylogenetic resolution. Finally, manual curation of large-scale databases such as SILVA is not feasible due to the growing number of sequences and frequent taxonomy updates (Glöckner et al., 2017).

Recent developments of sequencing technology and bioinformatic tools enabled generation of millions of high-quality full-length 16S rRNA reference sequences from any environmental ecosystem (Callahan et al., 2019; Karst et al., 2019, 2018). Obtaining full-length 16S rRNA gene sequences directly from environmental samples has made it possible to establish a near-complete reference database for high-taxonomic resolution studies of the wastewater treatment ecosystem, based on approximately one million 16S rRNA gene sequences retrieved directly from the wastewater treatment plants, which provided more than 9,500 exact sequence variants (ESVs) after dereplication and error-correction. Moreover, we created the AutoTax pipeline for generating a comprehensive ecosystem-specific taxonomic database with *de novo* names for novel taxa. The names are first inherited from sequences present in large-scale SILVA taxonomy, and *de novo* names are provided for all remaining unclassified taxa based on reproducible clustering with rank-specific identity thresholds (Yarza et al., 2014). These *de novo* names provide placeholder names for all novel phylotypes and act as fixed unique identifiers, making the taxonomic assignment independent of the analyzed dataset and enabling cross-study comparisons.

Here we present the MiDAS 3 reference database, which is based on the ESVs previously obtained from activated sludge and anaerobic digester systems (Dueholm et al., 2019), amended with additional sequences of potential importance in the field. These sequences represent bacteria from influent wastewater, potential pathogens, and genome-derived sequences of microbes found in wastewater treatment systems. The MiDAS 3 taxonomy is built using AutoTax (Dueholm et al., 2019) and proposes unique provisional names for all microorganisms important in wastewater treatment ecosystems at species-level resolution. We have furthermore curated some of these species-level names, and here we demonstrate the resolution power of MiDAS 3 by analyzing species-level community composition in 20 Danish full-scale WWTPs carrying out nutrient removal with chemical phosphorus removal (BNR) or enhanced biological phosphorus removal (EBPR) over 12 years. We provide important information about diversity, core communities, and the most abundant species in these ecosystems. Although the plants investigated are Danish, the results and approach are relevant for similar systems worldwide.

We also present the updated MiDAS Field Guide, an online platform that links the MiDAS 3 taxonomy-derived identity of all microbes from wastewater treatment systems to abundance information based on our long-term survey, and functional information (where available). MiDAS 3 alleviates important problems related to the taxonomic classification of microbes in environmental ecosystems, such as missing reference sequences and low taxonomic resolution, while our MiDAS Field Guide acts as an online knowledge repository facilitating understanding the function of microbes present in wastewater treatment ecosystems.

## Materials and methods

### MiDAS 3 reference database and taxonomy

The MiDAS 3 reference database was built using full-length 16S rRNA gene ESVs from 21 activated sludge WWTPs and 16 ADs systems, and taxonomic classification was assigned as described by Dueholm et al., (2019) and shown in **Figure 1**. The database was amended with 95 additional sequences from published genomes for the bacteria that are potentially important in wastewater treatment systems, but not present in the reference ESVs obtained (**Table S2**). Taxonomy was assigned to all sequences using the workflow presented in **Figure 1**. Gaps in the SILVA-derived taxonomy were filled with a *de novo* taxonomy that provides ‘midas_*x*_*y’* placeholder names, where *x* represents the first letter of taxonomic level, for which *de novo* assignment was created, and *y* is a number derived from the ESV that forms the cluster centroid for a given taxon. Where applicable, taxonomic classifications were curated with taxon names for ESVs that represent genomes published in peer-reviewed literature. Additionally, several manual curations were made, based on MiDAS 2.1 database and published literature (full list of amendments is presented in **Table S3**). MiDAS taxonomy file is available for download from the web platform (midasfieldguide.org/Downloads).

**Figure 1.**
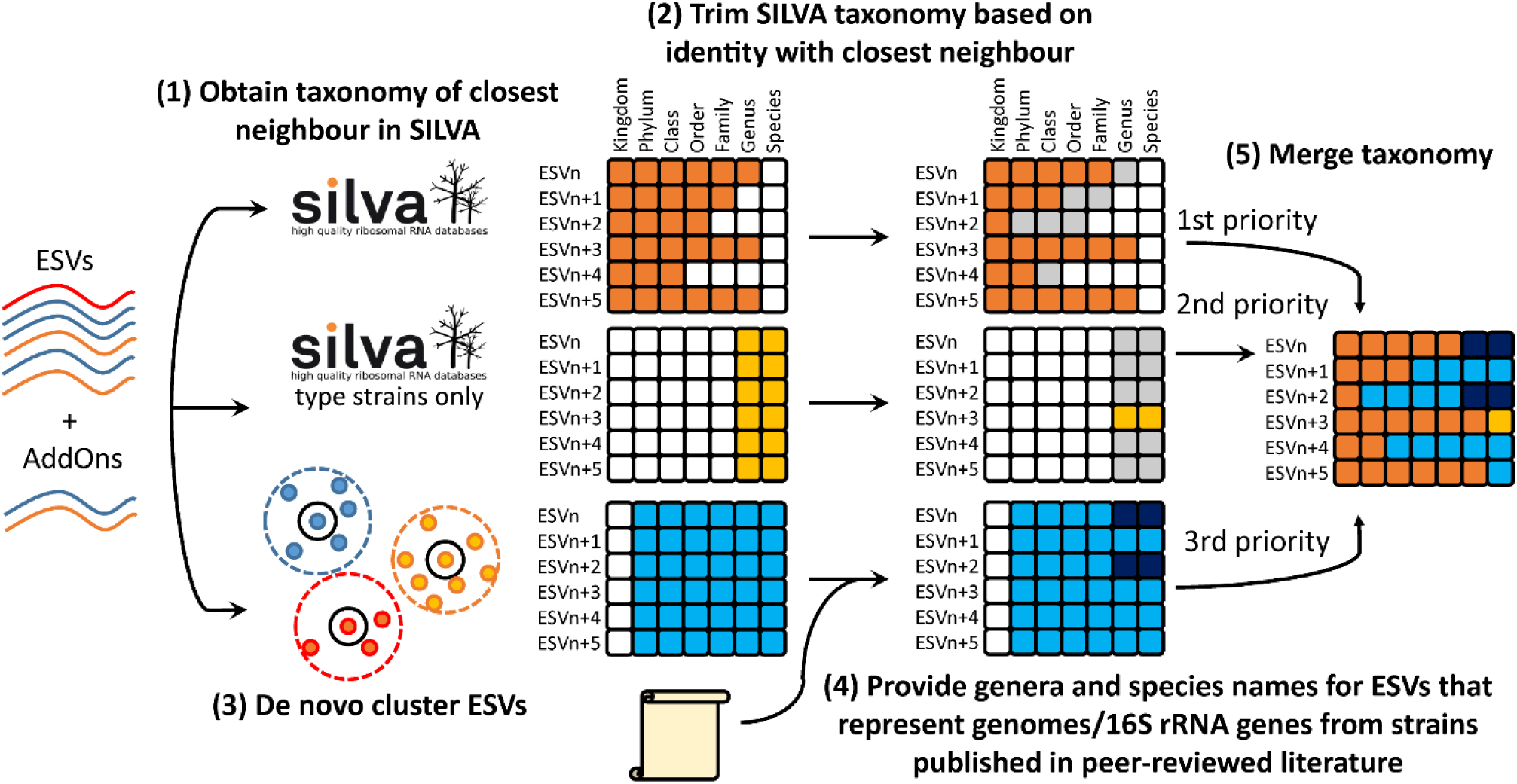
MiDAS 3 taxonomic assignment workflow based on the AutoTax framework. (1) ESVs from the study of Danish plants (Dueholm et al., 2019), amended with AddOns ESVs coming from, e.g., published genomes, are first mapped to the SILVA_132_SSURef_Nr99 database to identify the closest relative. (2) Taxonomy is adopted from this sequence after trimming, based on identity and the taxonomic thresholds proposed by Yarza et al., (2014) with 94.5% sequence similarity for genus-level. To gain species information, ESVs are mapped to sequences from type strains extracted from the SILVA database, and species names were adopted if the identity was >98.7% and the type strain genus matched that of the closest reference in the complete database. (3) ESVs were also clustered by greedy clustering at different identities, corresponding to the thresholds proposed by Yarza et al., (2014) to generate a stable *de novo* taxonomy. (4) The *de novo* taxonomy was curated with taxon names for ESVs that represent, e.g., genomes from strains published in peer-reviewed literature. (5) Finally, a comprehensive taxonomy was obtained by filling gaps in the SILVA-based taxonomy with the *de novo* taxonomy with provisional “midas” placeholder names (adapted from Dueholm et al., 2019).

### Activated sludge samples

Sampling of the activated sludge biomass from 20 Danish full-scale municipal WWTPs was carried out within the MiDAS project (McIlroy et al., 2015). The plants were sampled up to four times a year (February, May, August, October) from 2006 to 2018 with a total of 712 samples. All samples were taken from the aeration tank and sent overnight to Aalborg University for processing. Basic information about the design and operation of the WWTPs as well as number of samples collected are given in **Table S1**.

### Amplicon sequencing and bioinformatic analysis

DNA extraction, sample preparation including amplification of V1-3 region of 16S rRNA gene, and amplicon sequencing were conducted as described by Stokholm-Bjerregaard et al., (2017). Forward reads were processed as described by Dueholm et al., (2019) with raw reads trimmed to 250 bp, quality filtered, exact amplicon sequence variants (ASVs) identified, and taxonomy assigned using the MiDAS 3 reference database using the SINTAX classifier (Edgar, 2016).

### Data analysis and visualization

Data was imported into R (R Core Team, 2018) using RStudio (RStudio Team, 2015), analyzed and visualized using ggplot2 v.3.1.0 (Wickham, 2009) and ampvis2 v.2.4.9 (K. S. Andersen et al., 2018). The dataset containing 75,017 ASVs was rarefied to 10,000 reads per sample, resulting in 57,902 ASVs. Simpson’s alpha-diversity index calculation and redundancy analyses were performed using the ampvis2 package. Redundancy Analysis (RDA) was performed using the Hellinger transformation and constrained to WWTP location. For the analysis of rank abundance and core microbiome, all ASVs without taxonomic classification at the rank analyzed were removed. The remaining datasets consisting of 29,493 ASVs (50.9%) and 18,452 ASVs (31.9%) with genus- and species-level classification were normalized to 100%. These ASVs were represented in total by 1647 genera and 3651 species. The unclassified ASVs were primarily due to missing ESV reference sequences for rare ASVs, and the inability of the amplicons to resolve the species-level taxonomy for certain taxa. An example of the latter is genus *Trichococcus*, where the species *T. pasteurii, T. flocculiformis*, and *T. patagoniensis* cannot be distinguished even with full-length 16S rRNA gene sequences.

## Results and discussion

### The MiDAS reference database provides species-level classification

In the creation of the MiDAS 3 database, we have applied a novel approach (**Figure 1**), where high-throughput sequencing of high-quality full-length 16S rRNA genes (Karst et al., 2018) was combined with a new automatic taxonomy concept, AutoTax (Dueholm et al., 2019). The database contains full-length sequences from 21 Danish WWTPs carrying out nutrient removal, and 16 mesophilic and thermophilic Danish ADs. The MiDAS 3 reference database was additionally supplemented with high-quality full-length 16S rRNA gene sequences of microbes with known importance in the field, such as bacteria present in the influent wastewater and pathogens (**Table S2**).

MiDAS 3 taxonomic assignment is based on the AutoTax framework, where sequences first inherit taxonomic classification from the SILVA database (release 132, Quast et al., 2013), including species names (based on type strains), if available. The remaining unclassified sequences are assigned unique MiDAS names down to species level. Thus, the new release of MiDAS 3 taxonomy proposes a provisional genus and species name for all microorganisms found in activated sludge and anaerobic digester ecosystems, acting as stable identifiers and placeholder names for the sequences, until the representative microorganisms are further characterized and assigned approved names. Additionally, names derived from recently published genomes and other sources not yet incorporated in SILVA taxonomy, were manually curated (see **Table S3** for full list of curated names) to provide a near-complete database and taxonomy for microorganisms in wastewater treatment ecosystems. For a few species there were inconsistencies between the genus assigned by SILVA and the type strain name (see **Table S4**); these issues relate to the fact the SILVA taxonomy is updated based on phylogenetic analyses of the available 16S rRNA gene sequences, whereas type strain names have to be approved by the international taxonomy committee. The possibility to include additional sequences ensures that the database can easily be updated based on the current state of the ecosystem-specific field, and will drive the future releases of the MiDAS database.

Two examples of improved taxonomy are shown in **Table 1**. *Acidovorax* is one of the most abundant genera in Danish influent wastewater (McIlroy et al., 2017), with two abundant species identified using the MiDAS 3 reference database. *A. defluvii* is described by the correct classification down to genus level in all but one of the reference databases tested (MiDAS 2, SILVA, Greengenes, RDP). However, the other abundant species (midas_s_8989) was classified only at family level, with genus name either missing or assigned incorrectly by the other databases.

**Table 1.** Classification of abundant species belonging to genus *Acidovorax* and *Ca*. Amarolinea with different taxonomies.

| Taxonomy | Phylum | Class | Order | Family | Genus | Species |
| --- | --- | --- | --- | --- | --- | --- |
| **MiDAS 3** | Proteobacteria | Gammaproteobacteria | Betaproteobacteriales | Burkholderiaceae | *Acidovorax* | *A. defluvii* |
| MiDAS 2.1 | Proteobacteria | Betaproteobacteria | Burkholderiales | Comamonadaceae | *Acidovorax* | - |
| SILVA | Proteobacteria | Gammaproteobacteria | Betaproteobacteriales | Burkholderiaceae | *Acidovorax* | - |
| Greengenes | Proteobacteria | Betaproteobacteria | Burkholderiales | Comamonadaceae | - | - |
| RDP | Proteobacteria | Betaproteobacteria | Burkholderiales | Comamonadaceae | *Acidovorax* | - |
| **MiDAS 3** | Proteobacteria | Gammaproteobacteria | Betaproteobacteriales | Burkholderiaceae | *Acidovorax* | midas_s_8989 |
| MiDAS 2.1 | Proteobacteria | Betaproteobacteria | Burkholderiales | Comamonadaceae | - | - |
| SILVA | Proteobacteria | Gammaproteobacteria | Betaproteobacteriales | Burkholderiaceae | - | - |
| Greengenes | Proteobacteria | Betaproteobacteria | Burkholderiales | Comamonadaceae | - | - |
| RDP | Proteobacteria | Betaproteobacteria | Burkholderiales | Comamonadaceae | Simplicispira | - |
| **MiDAS 3** | Chloroflexi | Anaerolineae | Caldilineales | Amarolineaceae | *Ca. Amarolinea* | midas_s_1 |
| MiDAS 2.1 | Chloroflexi | SJA-15 | C10_SB1A | C10_SB1A | *Ca. Amarilinum* | - |
| SILVA | Chloroflexi | Anaerolineae | C10_SB1A | - | - | - |
| Greengenes | Chloroflexi | Anaerolineae | SHA-20 | - | - | - |
| RDP | - | - | - | - | - | - |
| **MiDAS 3** | Chloroflexi | Anaerolineae | Caldilineales | Amarolineaceae | *Ca. Amarolinea* | midas_s_553 |
| MiDAS 2.1 | Chloroflexi | SJA-15 | C10_SB1A | C10_SB1A | *Ca. Amarilinum* | - |
| SILVA | Chloroflexi | Anaerolineae | C10_SB1A | - | - | - |
| Greengenes | Chloroflexi | Anaerolineae | SHA-20 | - | - | - |
| RDP | - | - | - | - | - | - |
| **MiDAS 3** | Chloroflexi | Anaerolineae | Caldilineales | Amarolineaceae | *Ca. Amarolinea* | *Ca. A. aalborgensis* |
| MiDAS 2.1 | Chloroflexi | SJA-15 | C10_SB1A | C10_SB1A | *Ca. Amarilinum* | - |
| SILVA | Chloroflexi | Anaerolineae | C10_SB1A | - | - | - |
| Greengenes | Chloroflexi | Anaerolineae | SHA-20 | - | - | - |
| RDP | - | - | - | - | - | - |

Genus *Ca.* Amarolinea contains abundant filamentous bacteria in Danish WWTPs (Nierychlo et al., 2019) and can be associated with bulking incidents (M. H. Andersen et al., 2018; Nierychlo and Nielsen, 2017). All three abundant species belonging to the genus lack taxonomic classification below order level in public large-scale databases, preventing identification and analysis of this important member of the activated sludge system.

### Microbiome composition of Danish activated sludge plants

In this study, we performed a 12-year survey of bacteria present in 20 Danish full-scale WWTPs with nutrient removal that primarily treated municipal wastewater. All plants had biological nitrogen removal and most also had EBPR (17 plants), while 3 plants had chemical P-removal (BNR plants). The survey was carried out using 16S rRNA gene sequencing of the V1-3 region on the Illumina MiSeq platform. Using the MiDAS 3 taxonomy, microbial community composition was characterized at genus and species level in all plants based on 712 samples collected 2 to 4 times per year for each plant from 2006 to 2018.

The overall microbial diversity within each plant was estimated using the Simpson index of diversity and compared for all plants analyzed (**Figure 2**). The Simpson index places a greater weight on species evenness than the richness (Kim et al., 2017), thus allowing the assessment of how evenly distributed relative taxa contributions are across the samples. Since the value of the index represents the probability that two randomly selected individuals will belong to different taxa, the low values of Simpson index of diversity (**Figure 2**) indicate that the microbial communities were dominated by a low number of abundant taxa across the plants. Additionally, the low spread of the index in each plant indicates that the microbial communities across the plants were consistently stable across the years. Similar alpha diversity indices were observed in WWTPs across the world by Wu et al., (2019), calculated as Inverse Simpson (**Figure S1**).

**Figure 2.**
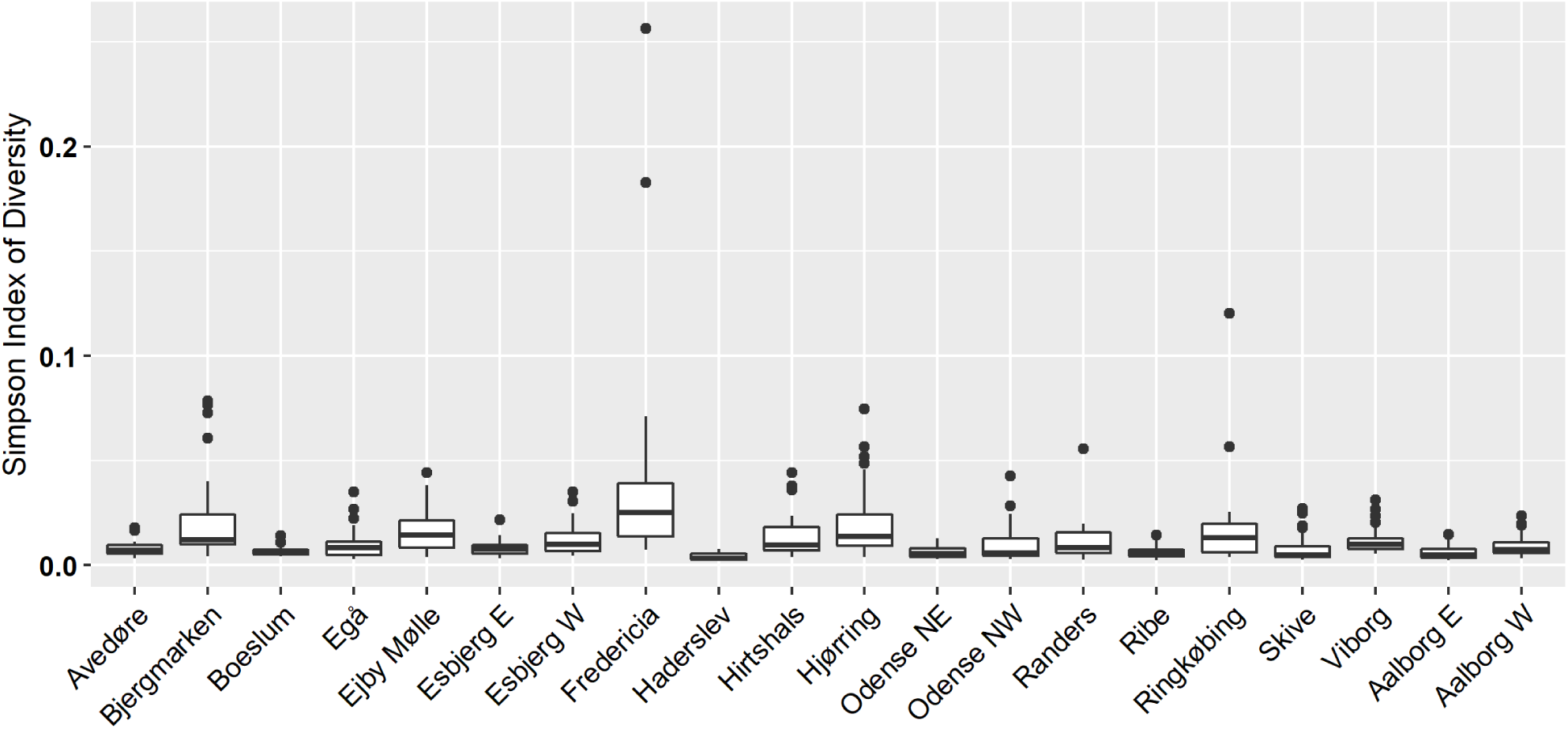
Alpha diversity estimates of activated sludge community composition using ASV-level taxa in individual plants, calculated using the Simpson index of diversity (1-D). Data represents 712 samples from 20 Danish full-scale WWTPs collected from 2006 to 2018 (with 17-51 samples per plant).

The overall variation in activated sludge community structure in 20 Danish WWTPs was visualized using RDA plot (**Figure 3**) with the WWTP location as the constrained variable. In many cases the samples coming from different WWTPs were not completely separated from each other, indicating high similarity between the microbial communities in those WWTPs. This is also highlighted by the fact that the eigenvalues of the two principal components plotted relative to the total variance in the data are relatively small (9.7% and 5.6%). Fredericia, Hirtshals, Esbjerg E, and Esbjerg W WWTPs appeared to have a more distinct microbial community composition, presumably due to a high fraction of industrial waste in the influent (**Table S1**). A pronounced clustering of the samples coming from individual WWTPs indicates that variation between the plants was likely more pronounced than within individual plants over time, and suggests long-term stability of the microbial communities in Danish plants.

**Figure 3.**
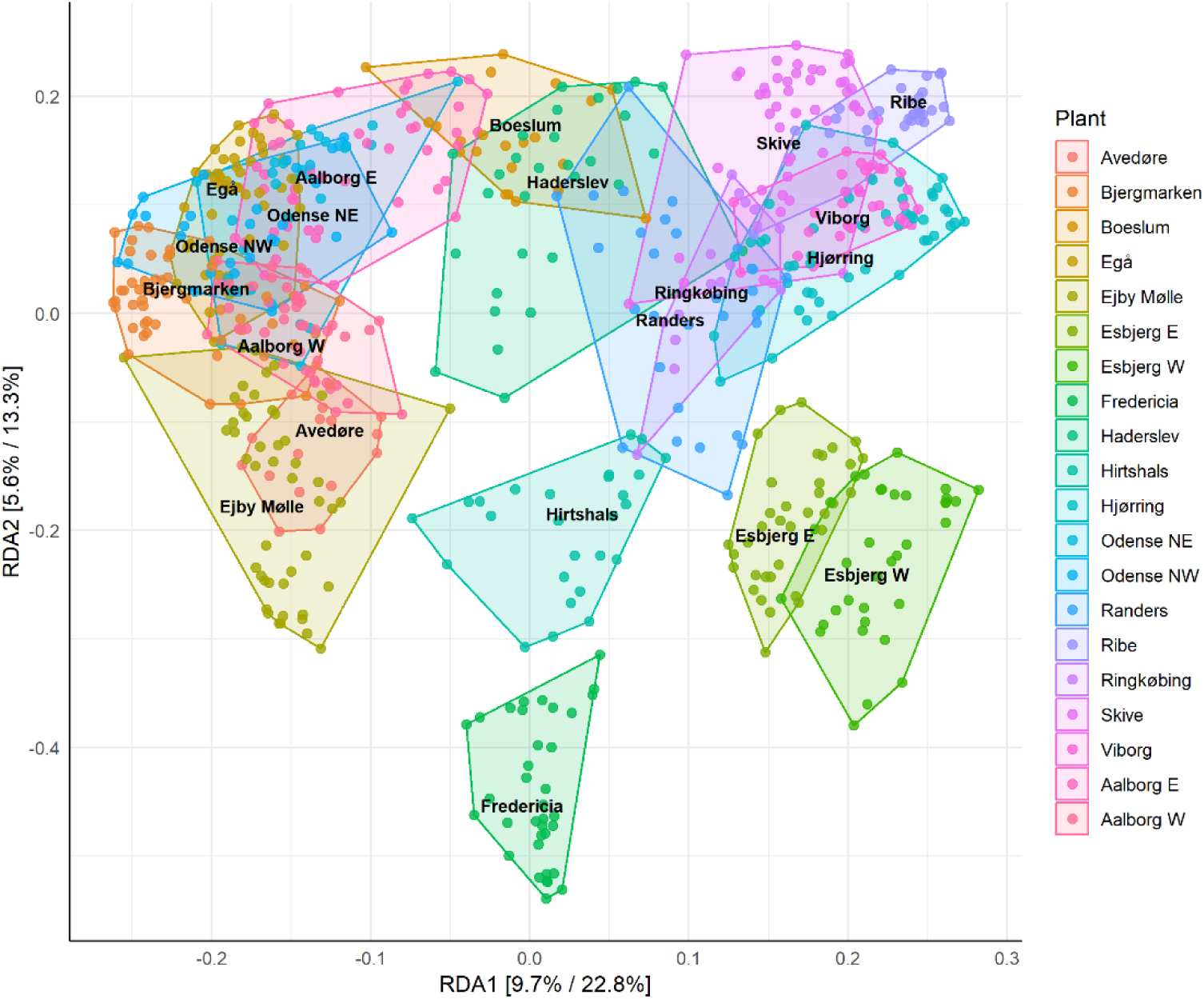
Redundancy analysis (RDA) plots of activated sludge communities, constrained to visualize the variance among 712 samples explained by WWTP location. Axes show the percent of total variance described by each component (first and second value represent relative contribution of each axis to the total variance in the data and the constrained space only, respectively). Data represents samples from 20 Danish full-scale WWTPs collected from 2006 to 2018.

The microbial diversity of the WWTPs was composed of 1,647 genera and 3,651 species. However, a relatively small fraction of these comprised a large proportion of the reads, thus representing the abundant taxa that are assumed to carry out the main functions in the system (**Figure 4a**) (Saunders et al., 2016). The abundant taxa were defined as those that constitute the top 80% of reads obtained by amplicon sequencing. In the BNR and EBPR plants, we found 157 and 164 abundant genera and 421 and 504 abundant species, respectively. These species had a relative ASV read abundance above approx. 0.04%.

**Figure 4.**
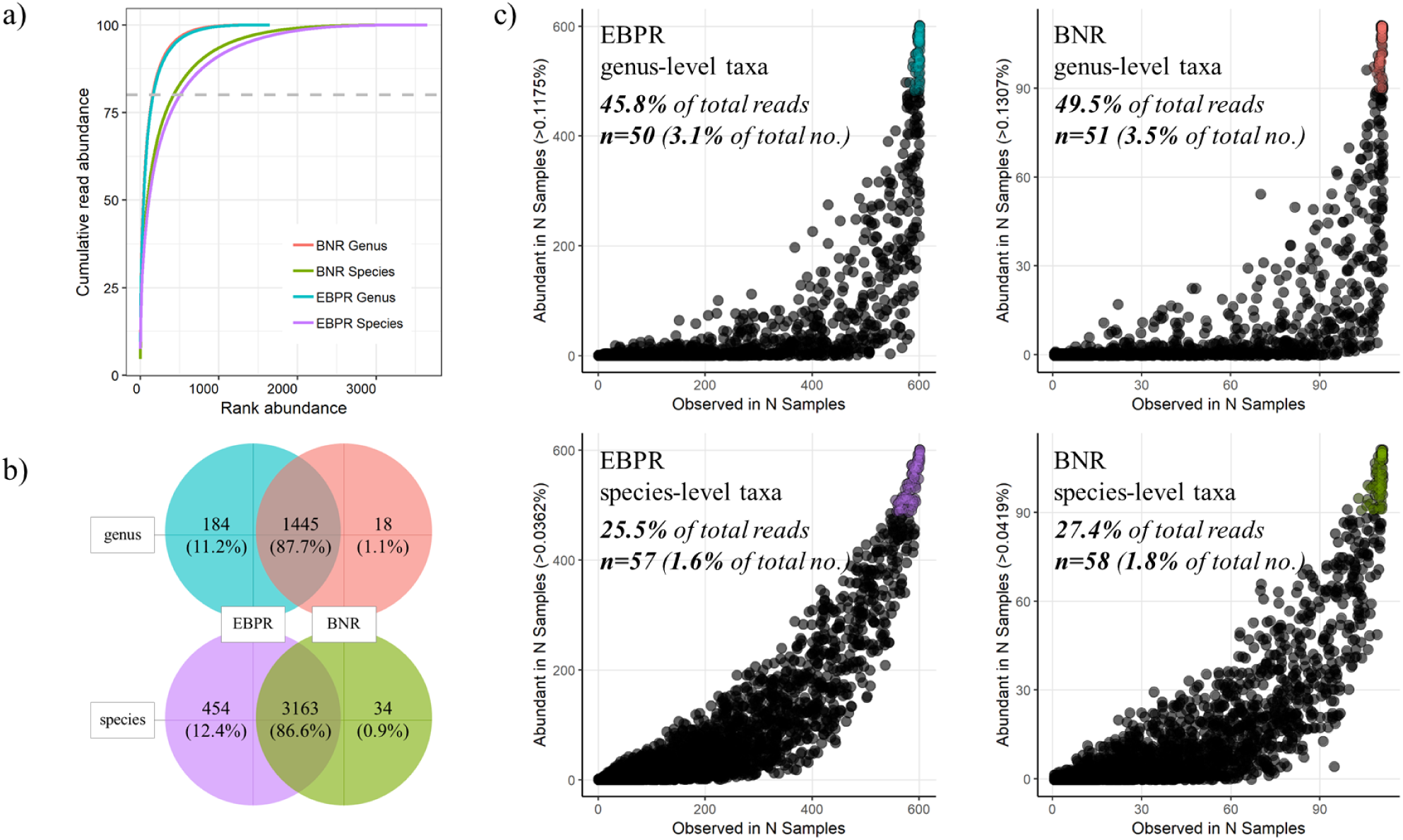
**a-c**. a) Cumulative read abundance of genus- and species-level taxa found in activated sludge, plotted in rank order. Dashed line denotes 80% of the reads, which is assumed to cover the abundant taxa, b) Venn-diagram showing abundant (top 80% of the reads) genera and species shared between BNR and EBPR plants, c) core community (observation frequency vs abundance frequency) in Danish BNR and EBPR plants for genus- and species-level taxa. Abundance cut-off values represent the lowest abundance of the taxon belonging to the top 80% of the reads in given group (genus/species). Darker points indicate multiple observations at given positions. Colored points designate core taxa. Data represents 111 samples from BNR plants and 601 samples from EBPR plants.

The overlap of taxa between the BNR and EBPR abundant communities both at genus and species level was high (**Figure 4b**), and the core taxa, defined as being present and abundant in ≥80% of the samples (Saunders et al., 2016), constituted more than 45% of the total reads at genus level and more than 25% of total reads at species level in both types of plants (**Figure 4c**). The number of genera and species belonging to the core communities in BNR and EBPR plants constituted a small fraction of the total number of taxa present (1-6-3.5%). Both types of plants shared 72% of genera and 65% of species in their respective core communities. The results show that the inclusion of an anaerobic process step in the EBPR plants compared to the conventional BNR plants did not affect the communities very much.

### Species-level community composition

The most abundant species present in the BNR and EBPR plants are shown in **Figure 5**. Among the 10 most abundant species, two belong to genera with a provisional ‘midas’ name, while of the top 50 and top 100 species, 13 and 36 species, respectively, belong to genera not possessing genus-level classification in SILVA (top 100 species are presented in **Figure S2**). These would be excluded from the analysis if MiDAS 3 was not used as the reference database, demonstrating the advantage of using the ecosystem-specific taxonomy.

**Figure 5.**
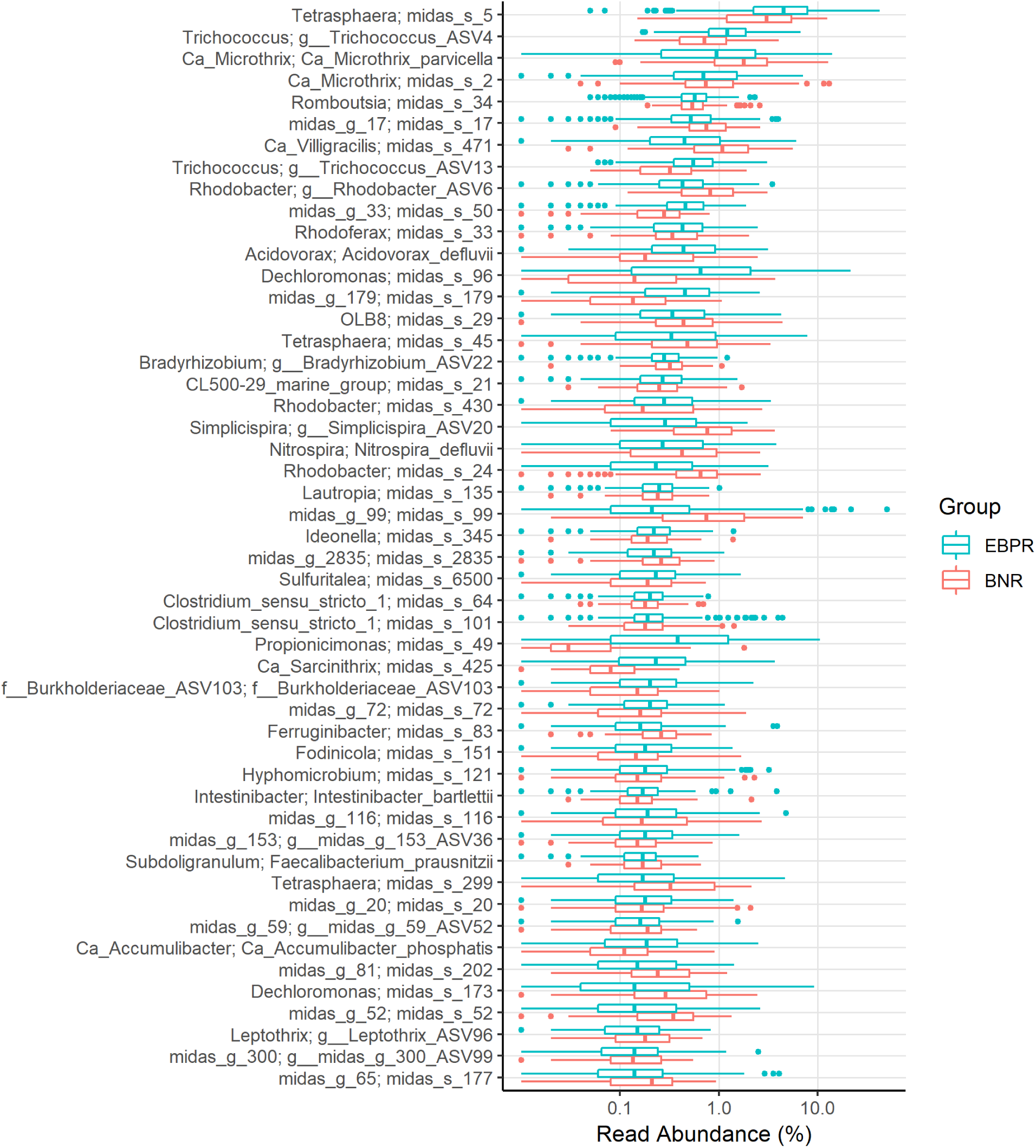
Boxplot showing the occurrence of the top 50 most abundant species in Danish BNR and EBPR WWTPs.

A large fraction of the most abundant species is poorly described in the literature. The top 50 species belong to 41 different genera, of which almost 60% (28 genera) have no known function in wastewater treatment systems (McIlroy et al., 2017, 2015). The function of 65% of all genera defined as abundant in the EBPR and BNR plants (**Figure 2a**) is unknown, demonstrating the need for more studies that link identity and function. A few species, however, match SILVA type strains, such as *Acidovorax defluvii* and *Intestinibacter bartletti*, so these may be applied for deeper studies of their physiology.

The most abundant species in both BNR and EBPR plants belong to the genus *Tetrasphaera,* which represents polyphosphate-accumulating organisms (PAO) with versatile metabolism, including the potential for denitrification (Herbst et al., 2019; Kristiansen et al., 2013). Other very abundant species belong to the filamentous genera *Ca.* Microthrix (McIlroy et al., 2013) and *Ca*. Villigracilis (Nierychlo et al., 2019); the former known to cause serious bulking problems (Rossetti et al., 2005). *Trichococcus* may also possess filamentous morphology in activated sludge and may be implicated in bulking (Liu and Seviour, 2001). *Rhodoferax* is a denitrifier (McIlroy et al., 2014), while the function in activated sludge is unknown for species in the *Romboutsia* and *Rhodobacter* genera. The novel genus midas_g_17, for which no functional information is available, belongs to the family Saprospiraceae, and was wrongly classified as *Ca*. Epiflobacter in the previous version of MiDAS taxonomy. Based on the coverage of the FISH probes available and mapping of the partial 16S sequences (Xia et al., 2008) to MiDAS 3 database, *Ca*. Epiflobacter name was re-assigned to a correct genus, which is present at much lower abundance in the ecosystem. MiDAS 3 provides good opportunities to reclassify sequences from previous studies, for which information about distribution and/or function is available. For example, after mapping a set of partial 16S rRNA gene sequences originally related to genus *Aquaspirillum* (Thomsen et al., 2007), they could be reclassified as midas_g_33 – an abundant novel genus, indicating its potential role in denitrification.

Interestingly, the diversity of abundant species within the different genera was generally low in the plants, typical with 3-4 abundant species in each of the most abundant genera. However, the known PAO *Tetrasphaera* and the less abundant *Ca.* Accumulibacter exhibited slightly higher diversity, each having 10 and 5 abundant species, respectively (**Figure 6**). *Tetrasphaera* had one dominant species (midas_s_5) present in most plants, while the other species appeared more randomly distributed. No clear dominance of any of *Ca.* Accumulibacter species could be observed as their abundance was very different across the plants. Two of these, *Ca*. A. phosphatis and *Ca.* A. aalborgensis are described at the species-level, based on the genomes available (Albertsen et al., 2016; Martín et al., 2006). Interestingly, the genus *Dechloromonas*, which is also suggested to be a putative PAO (Stokholm-Bjerregaard et al., 2017), was more abundant than *Ca*. Accumulibacter, and their diversity was low with only two dominant species present in almost all plants. In general, high species diversity of the PAOs may suggest a high degree of specialization allowing the exploitation of different niches in the activated sludge ecosystem. *Ca.* Accumulimonas, another putative PAO, which was described in the previous version of MiDAS, is not included in the present taxonomy version, as sequences representing that genus, according to the present SILVA taxonomy, were classified to the genus *Halomonas*.

**Figure 6.**
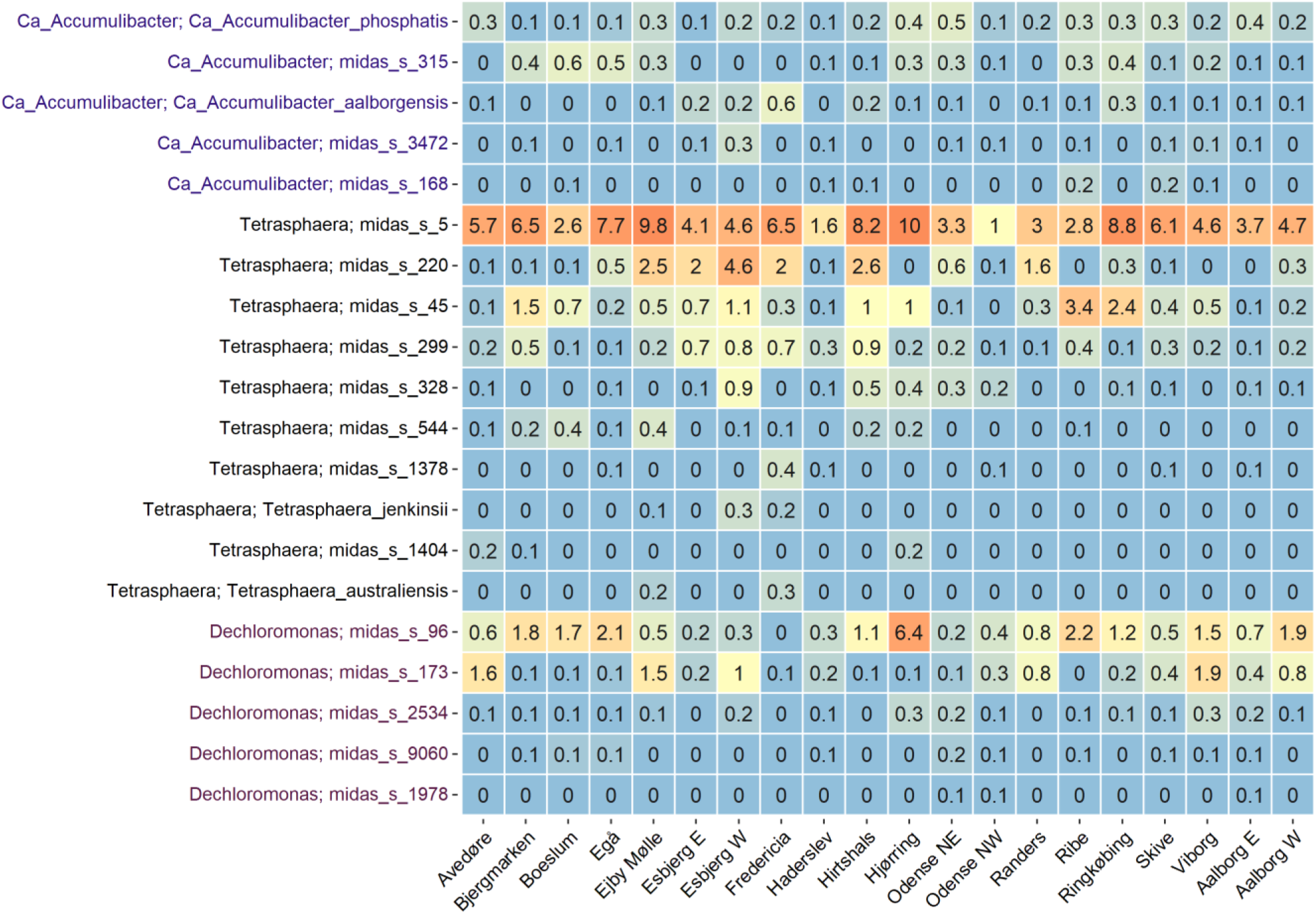
Occurrence of abundant PAO species in 20 full-scale activated sludge WWTPs. Data represents average read abundance values for each plant, based on samples collected 2–4 times per year from 2006 to 2018. Individual PAO genera are indicated by different colors.

### The MiDAS reference database provides a common language for the field

Only a few studies have made surveys of the microbial communities in activated sludge systems, and in all cases, it has only been with genus-level and not species-level classification (Saunders et al., 2016; Wu et al., 2019). Since the different studies have applied different extraction procedures, different primers (typically V1-3 or V4), and different databases for taxonomic assignment (MiDAS 2, SILVA, RDP or Greengenes), it can be difficult to compare results across studies at the genus level, and impossible at lower taxonomic levels. Therefore, the use of the ecosystem-specific MiDAS database will provide a common language for all work in the field. Another key advantage of the MiDAS 3 taxonomy is that it includes taxa abundant in similar systems across the world (Dueholm et al., 2019), which are missing in SILVA, RDP, and Greengenes. By using this approach, and similar protocols for extraction and primer selection (Albertsen et al., 2015; Dueholm et al., 2019), it will be possible to compare the results of WWTP microbiome studies in the future.

### The MiDAS reference database for anaerobic digesters

Anaerobic digestion is a key technology increasingly used worldwide to reduce waste, generate energy, and minimize the carbon footprint of the plant. In order to provide a comprehensive reference database for the microbes present in the whole wastewater treatment system, MiDAS 3 is based on sequences from activated sludge as well as 16 full-scale Danish anaerobic digesters treating primary and waste activated sludge. A detailed analysis of bacterial and archaeal community structure at species-level has been performed in another study (Jiang et al., n.d.). The study was based on >1000 samples collected over 6 years from 46 anaerobic digesters, located at 22 WWTPs in Denmark. When the microbial data was coupled with the analysis of operational, physicochemical, and performance parameters, the main factors driving the assembly of the AD community were elucidated.

### MiDAS Field Guide online platform (midasfieldguide.org)

The new MiDAS Field Guide (www.midasfieldguide.org) is a comprehensive database of microbes in wastewater treatment systems, which was updated at the same time as the release of the MiDAS 3 database took place. All microbes found in activated sludge and anaerobic digesters are now included in the field guide and, for the first time, species are listed for each genus. Our database now covers more than 1,800 genera and 4,200 species found in biological nutrient removal treatment plants and anaerobic digesters. More detailed abundance information is now provided both at genus and species level, and functional information is described where available.

The MiDAS Field Guide integrates the identity of the microbes with the available functional information (**Figure 7**) into a community knowledge platform in the form of a searchable database of referenced information, thus acting as a central, on-line repository for current knowledge, where all are welcome to contribute. Due to the rapidly growing amount of relevant literature, and high number of taxa included, the MiDAS Field Guide is a continuous work in progress, with our efforts prioritized towards the microorganisms that comprise the core microbiome as well as other abundant or process-critical species.

**Figure 7.**
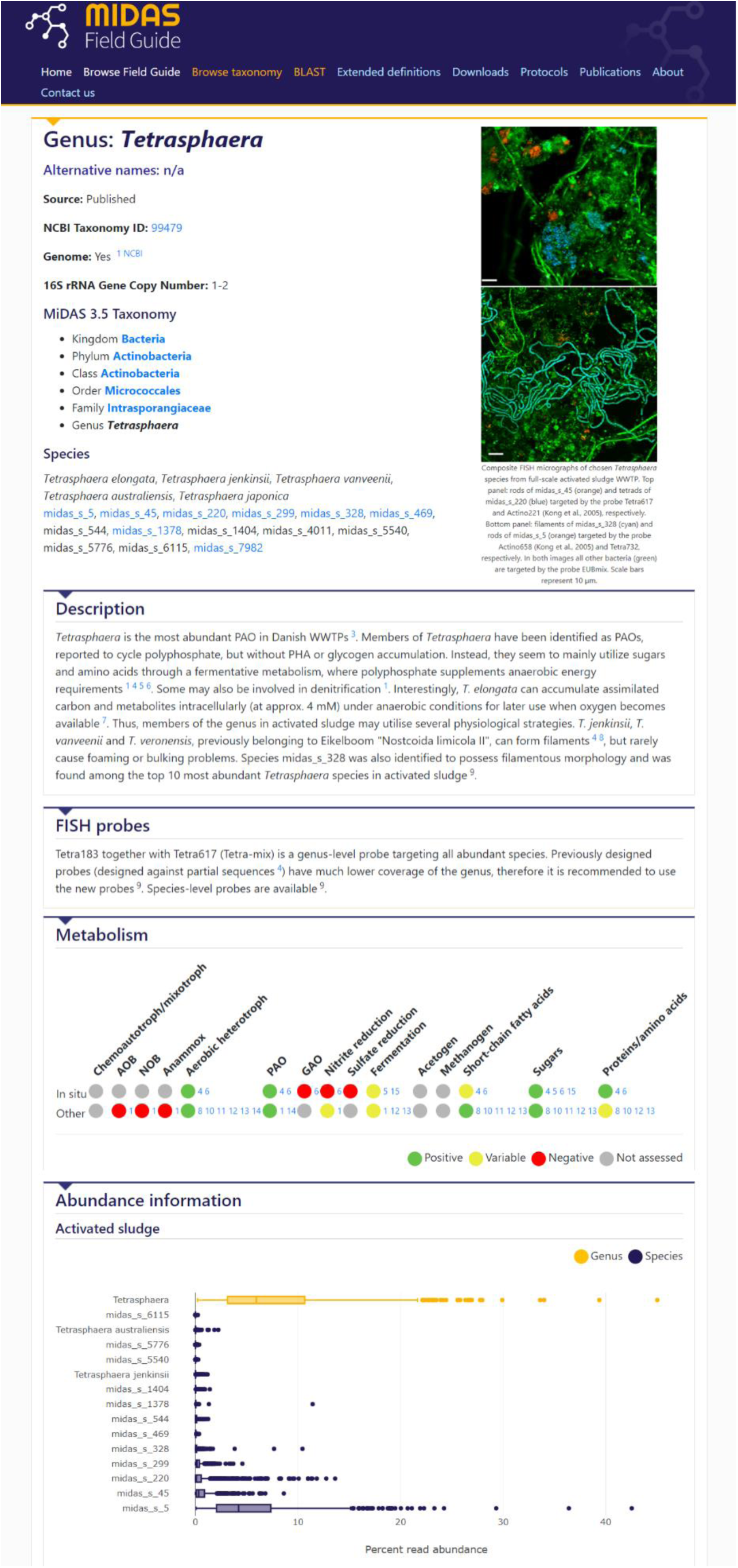
MiDAS Field Guide example of *Tetrasphaera* genus entry.

New features launched in the updated MiDAS Field Guide include: 1) phylogenetic *Search by taxonomy* function, where all microbes found in the ecosystem are displayed in their respective taxonomic groups, 2) the possibility of classifying sequences using blast function, and MiDAS 3 taxonomy directly online. Updated protocols for DNA extraction and amplicon library preparation are available online to facilitate establishment of common framework for studies of microbial ecology of wastewater treatment systems.

## Conclusions

Here we present the new ecosystem-specific MiDAS 3 reference database, based on a comprehensive set of full-length 16S rRNA gene sequences derived from activated sludge and anaerobic digester systems, and MiDAS 3 taxonomy built using AutoTax. It proposes unique provisional names for all microorganisms important in wastewater treatment ecosystems, down to species level. MiDAS 3 was used in the detailed analysis of microbial communities in 20 Danish wastewater treatment plants (WWTPs) with nutrient removal, sampled over 12 years. This enabled unprecedented resolution, revealing for the first time all species present in the plants, the stability of the communities, many abundant core taxa, and the diversity of species in important functional guilds, exemplified by the PAOs. We found many genera without known function, emphasizing the need for more efforts towards elucidating the role of important members of wastewater treatment ecosystems. We also present a new version of the MiDAS Field Guide, an online resource with a complete list of genera and species found in activated sludge and anaerobic digesters. Through the field guide, microbe identity is linked to available knowledge on function, and user microbial 16S rRNA gene sequences can be classified online with the MiDAS 3 taxonomy. The central purpose of the MiDAS Field Guide is to facilitate collaborative research into wastewater treatment ecosystem functions, which will support efforts towards the sustainable production of clean water and bioenergy, and the development of resource recovery, ending ultimately in the circular economy.

## Acknowledgement

The project has been funded by the Danish Research Council (grant 6111-00617A), Innovation Fund Denmark (OnlineDNA, grant 6155-00003A), the Villum foundation (Dark Matter and grant 13351), and Danish wastewater treatment plants.

