## Supplementary information for "Species-level microbiome composition of activated sludge - introducing the MiDAS 3 ecosystem-specific reference database and taxonomy"

### Supplementary Tables:

**Table S1.** Design, operation, and sample number from WWTPs included in the study.

**Table S2.** List of genome- or pure culture-derived sequences added to MiDAS 3 reference database.

**Table S3.** List of bacterial names curated in MiDAS 3 reference database, based on available genomes, published literature, and transferred from MiDAS 2.1 taxonomy.

**Table S4.** List of bacterial names with differences between genus name and type strain name inherited from Silva_132_SSURef_Nr99 (Quast et al., 2013).

### Supplementary Figures:

**Figure S1.** Alpha diversity estimate of activated sludge community composition using ASV-level taxa in individual plants, calculated using the Inverse Simpson index. Data represents 712 samples from 20 Danish full-scale WWTPs collected from 2006 to 2018 (with 17-51 samples per plant).

**Figure S2.** Boxplot showing the occurrence of top 100 most abundant species in Danish EBPR and BNR WWTPs.

**Table S1**. Design, operation, and sample number from WWTPs included in the study.

| WWTP name | PE | Configuration | Industrial waste (% of COD) | Primary settling | Design | EBPR configuration | Digester | No. of samples |
|---|---|---|---|---|---|---|---|---|
| Bjergmarken | 225 000 | Alternating | 20 | No | EBPR | MAT | Yes | 51 |
| Boeslum | 23 000 | Recirculation | 5 | No | EBPR | MAT | No | 22 |
| Egå | 120 000 | Alternating | 40 | Yes | EBPR | SSH | Yes | 51 |
| Ejby Mølle | 410 000 | Alternating | 49-55 | Yes | EBPR | MAT | Yes | 47 |
| Esbjerg E | 120 000 | Recirculation | 60 | Yes | BNR | - | Yes | 36 |
| Esbjerg W | 290 000 | Recirculation | 60 | Yes | BNR | - | Yes | 35 |
| Fredericia | 420 000 | Recirculation | 75 | No | EBPR | MAT | Yes | 36 |
| Haderslev | 100 000 | Alternating | 5 | No | EBPR | SSH | No | 30 |
| Hirtshals | 117 000 | Alternating | 60-70 | No | EBPR | MAT | No | 26 |
| Hjørring | 120 000 | Recirculation | 30 | No | EBPR | MAT | Yes | 49 |
| Odense NE | 36 000 | Alternating | 20 | No | EBPR | MAT | Yes | 30 |
| Odense NW | 75 000 | Alternating | 18 | No | BNR | - | Yes | 29 |
| Randers | 130 000 | Recirculation | 5 | Yes | EBPR | SSH | Yes | 30 |
| Ribe | 25 000 | Recirculation | 20 | No | EBPR | SSH | No | 32 |
| Ringkøbing | 42 500 | Alternating | 10 | Yes | EBPR | SSH | Yes | 17 |
| Skive | 123 000 | Recirculation | 20-65 | No | EBPR | SSH | No | 49 |
| Viborg | 80 000 | Recirculation | 10 | Yes | EBPR | SSH | Yes | 33 |
| Aalborg E | 150 000 | Alternating | 25 | Yes | EBPR | SSH | Yes | 48 |
| Aalborg W | 330 000 | Alternating | 25 | Yes | EBPR | SSH | Yes | 44 |
| Avedøre | 345 000 | Alternating | 25 | Yes | EBPR | MAT | Yes | 17 |

EBPR: enhanced biological P-removal; BNR biological removal of N and chemical removal of P; MAT: mainstream anaerobic tank; SSH: return sludge sidestream hydrolysis.

**Table S2.** List of genome- or pure culture-derived sequences added to MiDAS 3 reference database.

| No. | Add-on sequence name | Accession | Source |
|---|---|---|---|
| 1 | Aeromonas caviae strain FDAARGOS_72 | NZ_CP026055 | |
| 2 | Bacillus anthracis str. Ames | NC_003997 | PMID=12721629 |
| 3 | Brevefilum fermentans | NZ_LT859958 | PMID=28690595 |
| 4 | Campylobacter coli strain aerotolerant OR12 | NZ_CP019977 | |
| 5 | Campylobacter jejuni subsp. jejuni NCTC 11168 | NC_002163 | PMID=17565669 |
| 6 | candidate division SR1 bacterium Aalborg_AAW-1 | CP011268 | PMID=26067967 |
| 7 | Candidatus Accummulibacter aalborgensis | FLQX01000078 | PMID=27458436 |
| 8 | Candidatus Accumulibacter delftensis | | PMID=31189123 |
| 9 | Candidatus Accumulibacter phosphatis | CP001715 | PMID=16998472 |
| 10 | Candidatus Amarolinea aalborgensis | MH537630 | PMID=30146409 |
| 11 | Candidatus Bipolaricaulis anaerobius | AYTS01000061 | PMID=29884828 |
| 12 | Candidatus Brocadia caroliniensis | AYTS01000061 | PMID=28088723 |
| 13 | Candidatus Brocadia sapporoensis strain 40 | NZ_MJUW02000026 | PMID=27932661 |
| 14 | Candidatus Brocadia sinica JPN1 | NZ_BAFN01000001 | PMID=25883286 |
| 15 | Candidatus Competibacter denitrificans | NZ_CBTJ020000028 | PMID=24173461 |
| 16 | Candidatus Contendobacter odensis | CBTK010000065 | PMID=24173461 |
| 17 | Candidatus Defluviicoccus seviourii | UXAT01000031 | PMID=30476038 |
| 18 | Candidatus Fermentibacter daniensis | LKHB01000197 | PMID=27058503 |
| 19 | Candidatus Microthrix calida strain TNO | DQ147284 | PMID=16913916 |
| 20 | Candidatus Microthrix parvicella | CANL01000022 | PMID=23446830 |
| 21 | Candidatus Nitrospira defluvii | NC_014355 | PMID=20624973 |
| 22 | Candidatus Nitrospira inopinata | LN885086 | PMID=26610024 |
| 23 | Candidatus Nitrospira nitrosa | CZQA01000015 | PMID=26610025 |
| 24 | Candidatus Nitrotoga sp. KNB | LS423452 | PMID=29991589 |
| 25 | Candidatus Promineofilum breve | LN890655 | PMID=26905629 |
| 26 | Candidatus Propionivibrio aalborgensis | FLQY01000362 | PMID=27458436 |
| 27 | Candidatus Saccharimonas aalborgensis | CP005957 | PMID=23707974 |
| 28 | Dechloromonas denitrificans strain | NZ_LODL01000035 | ATCC BAA-841 |
| 29 | Enterobacter cloacae subsp. cloacae ATCC 13047 | NC_014121 | PMID=20207761 |
| 30 | Escherichia coli O157:H7 str. Sakai | NC_002695 | PMID=11258796 |
| 31 | Helicobacter pylori 26695 | NNC_000915 | PMID=9252185 |
| 32 | Methanobacterium bryantii | LMVM01000014 | PMID=28826405 |
| 33 | Methanobacterium formicicum DSM 3637 | NZ_AMPO01000020 | PMID=23209223 |
| 34 | Methanobacterium lacus strain AL-21 | NC_015216 | PMID=24449792 |
| 35 | Methanobacterium paludis strain SWAN1 | NC_015574 | PMID=24449792 |
| 36 | Methanobrevibacter boviskoreani | NZ_BAGX02000040 | PMID=23469331 |
| 37 | Methanobrevibacter olleyae strain YLM1 | NZ_CP014265 | PMID=27056228 |
| 38 | Methanobrevibacter ruminantium M1 | NC_013790 | PMID=20126622 |
| 39 | Methanobrevibacter smithii ATCC 35061 | NC_009515 | PMID=17563350 |
| 40 | Methanocaldococcus bathoardescens | NZ_CP009149 | PMID=25634941 |
| 41 | Methanocaldococcus jannaschii DSM 2661 | NC_000909 | PMID=8688087 |
| 42 | Methanocella arvoryzae MRE50 | NC_009464 | PMID=16857943 |
| 43 | Methanocella conradii HZ254 | NC_017034 | PMID=22493204 |
| 44 | Methanocella paludicola SANAE | NC_013665 | PMID=18398197 |
| 45 | Methanococcoides methylutens strain DSM 2657 | NZ_JRHO01000009 | PMID=25414501 |
| 46 | Methanocorpusculum labreanum | NC_008942 | PMID=21304657 |
| 47 | Methanocorpusculum parvum | NZ_LMVO01000026 | PMID=28826405 |
| 48 | Methanoculleus horonobensis strain T10 | NZ_BCNY01000003 | PMID=27034500 |

| | | | |
|---|---|---|---|
| 49 | Methanoculleus marisnigri JR1 | NC_009051 | PMID=21304656 |
| 50 | Methanoculleus sediminis | NZ_JXOJ01000002 | PMID=25855623 |
| 51 | Methanoculleus taiwanensis strain CYW4 | NZ_LHQS01000005 | PMID=25575827 |
| 52 | Methanoculleus thermophilus | NZ_BCNX01000018 | PMID=27034500 |
| 53 | Methanofollis ethanolicus strain HASU | NZ_BCNW01000001 | PMID=27034500 |
| 54 | Methanolacinia petrolearia DSM 11571 | NC_014507 | PMID=21304750 |
| 55 | Methanolinea tarda NOBI-1 | NZ_AGIY02000001 | PMID=25189585 |
| 56 | Methanolobus psychrophilus | NC_018876 | PMID=23760934 |
| 57 | Methanomassiliicoccus luminyensis B10 | NZ_CAJE01000013 | PMID=22887657 |
| 58 | Methanoregula formicica SMSP | NC_019943 | PMID=25189582 |
| 59 | Methanosarcina spelaei | NZ_LMVP01000249 | PMID=28826405 |
| 60 | Methanosphaera stadtmanae DSM 3091 | NC_007681 | PMID=16385054 |
| 61 | Methanosphaerula palustris E1-9c | NC_011832 | PMID=26543115 |
| 62 | Methanospirillum hungatei JF-1 | NC_007796 | PMID=26744606 |
| 63 | Methanothermobacter marburgensis | NC_014408 | PMID=20802048 |
| 64 | Methanothermobacter thermautotrophicus | NC_000916 | PMID=9371463 |
| 65 | Methanothermus fervidus DSM 2088 | NC_014658 | PMID=21304736 |
| 66 | Methanothrix harundinacea | NC_017527 | PMID=22590603 |
| 67 | Micropruina glycogenica | NZ_LT985188 | PMID=29875741 |
| 68 | Mycobacterium avium subsp. paratuberculosis | NC_002944 | PMID=16116077 |
| 69 | Mycobacterium intracellulare ATCC 13950 | NC_016946 | PMID=22535933 |
| 70 | Mycobacterium tuberculosis H37Rv | NC_000962 | PMID=20980199 |
| 71 | Neomegalonema perideroedes | NZ_KB893658 | PMID=26203335 |
| 72 | Nitrobacter hamburgensis X14 | NC_007964 | PMID=18326675 |
| 73 | Nitrobacter winogradskyi | NC_007406 | PMID=16517654 |
| 74 | Nitrosomonas communis strain Nm2 | NZ_CP011451 | PMID=26769932 |
| 75 | Nitrosomonas europaea ATCC 19718 | NC_004757 | PMID=12700255 |
| 76 | Nitrosomonas eutropha C91 | NC_008344 | PMID=17991028 |
| 77 | Nitrospira moscoviensis | CP011801 | PMID=26305944 |
| 78 | Pseudomonas aeruginosa PAO1 | NC_002516 | PMID=18978025 |
| 79 | Pseudomonas monteilii | CP006978 | PMID=24874689 |
| 80 | Pyrococcus furiosus DSM 3638 | NC_003413 | PMID=11210495 |
| 81 | Rhodococcus pyridinivorans | CP006996 | PMID=24874690 |
| 82 | Salmonella enterica subsp. enterica serovar Typhimurium str. LT2 | NC_003197 | PMID=11677609 |
| 83 | Shigella boydii Sb227 | NC_007613 | PMID=16275786 |
| 84 | Shigella dysenteriae Sd197 | NC_007606 | PMID=16275786 |
| 85 | Shigella flexneri 2a str. 301 | NC_004337 | PMID=12384590 |
| 86 | Shigella sonnei strain FDAARGOS_90 | NZ_CP014099 | |
| 87 | Staphylococcus aureus subsp. aureus NCTC 8325 | NC_007795 | |
| 88 | Tetrasphaera australiensis Ben110 | NZ_HG764815 | PMID=23178666 |
| 89 | Tetrasphaera elongata Lp2 | NZ_HF570956 | PMID=23178666 |
| 90 | Tetrasphaera japonica T1-X7 | NZ_HF570958 | PMID=23178666 |
| 91 | Tetrasphaera jenkinsii Ben 74 | NZ_HF571038 | PMID=23178666 |
| 92 | Thiothrix eikelboomii strain | NZ_FUYB01000034 | ATCC 49788 |
| 93 | Vibrio cholerae O1 biovar El Tor str. N16961 | NC_002505 | PMID=10952301 |
| 94 | Vibrio parahaemolyticus RIMD 2210633 | NC_004603 | PMID=12620739 |
| 95 | Yersinia enterocolitica subsp. enterocolitica 8081 | NC_008800 | PMID=17173484 |

**Table S3.** List of bacterial names curated in MiDAS 3 reference database, based on available genomes, published literature, and transferred from MiDAS 2.1 taxonomy.

| Previous name | MiDAS 3 name | Support | Reference |
|---|---|---|---|
| Acetothermia | Ca_Bipolaricaulota | genome available | PMID=29884828 |
| Acetothermiia | Ca_Bipolaricaulia | genome available | PMID=29884828 |
| ADurb.Bin120 | Ca_Brevefilum | genome available | PMID=28690595 |
| Aegiribacteria | Ca_Fermentibacterota | first published genome | PMID=27058503 |
| C10-SB1A | Caldilineales | genome available | PMID=30146409 |
| Ca_Nitrotoga | Nitrotoga | has been isolated | PMID=29991589 |
| Meganema | Neomegalonema | has been renamed | PMID=26203335 |
| Meganema_perideroedes | Neomegalonema_perideroedes | has been renamed | PMID=26203335 |
| Methanosaeta | Methanothrix | has been renamed | doi: 10.1099/ijs.0.037366-0 |
| Methanosaeta_concilii | Methanothrix_concilii | has been renamed | doi: 10.1099/ijs.0.037366-0 |
| Methanosaeta_harundinacea | Methanothrix_harundinacea | has been renamed | doi: 10.1099/ijs.0.037366-0 |
| Selenomonadales | Acidaminococcales | has been renamed | PMID=25999592 |
| midas_s_368 | Ca_Accumulibacter_phosphatis | genome available | PMID=16998472 |
| midas_s_2676 | Ca_Accumulibacter_aalborgensis | genome available | PMID=27458436 |
| midas_f_1 | Amarolineaceae | genome available | PMID=30146409 |
| midas_g_1 | Ca_Amarolinea | genome available | PMID=30146409 |
| midas_s_2372 | Ca_Amarolinea_aalborgensis | genome available | PMID=30146409 |
| midas_f_9589 | Ca_Bipolaricaulaceae | genome available | PMID=29884828 |
| midas_o_9589 | Ca_Bipolaricaulales | genome available | PMID=29884828 |
| midas_g_9589 | Ca_Bipolaricaulis | genome available | PMID=29884828 |
| midas_s_9589 | Ca_Bipolaricaulis_anaerobius | genome available | PMID=29884828 |
| midas_s_234 | Ca_Brevefilum_fermentans | genome available | PMID=28690595 |
| midas_s_9602 | Ca_Brocadia_caroliniensis | genome available | PMID=28088723 |
| midas_s_9608 | Ca_Brocadia_sapporoensis | genome available | PMID=27932661 |
| midas_s_9609 | Ca_Brocadia_sinica | genome available | PMID=25883286 |
| midas_s_127 | Ca_Competibacter_denitrificans | genome available | PMID=24173461 |
| midas_s_866 | Ca_Contendobacter_odensis | genome available | PMID=24173461 |
| midas_s_1030 | Ca_Defluviicoccus_seviourii | genome available | PMID=30476038 |
| midas_g_23 | Ca_Epiflobacter | manual mapping of 35 partial 16S seqs from original publication | PMID=18263744 |
| midas_g_18 | Ca_Fermentibacter | genome available | PMID=27058503 |
| midas_s_18 | Ca_Fermentibacter_daniensis | genome available | PMID=27058503 |
| midas_f_18 | Ca_Fermentibacteraceae | genome available | PMID=27058503 |
| midas_o_18 | Ca_Fermentibacterales | genome available | PMID=27058503 |
| midas_c_18 | Ca_Fermentibacteria | genome available | PMID=27058503 |
| midas_s_9576 | Ca_Microthrix_calida | has been isolated | PMID=16913916 |
| midas_s_3 | Ca_Microthrix_parvicella | genome available | PMID=23446830 |
| midas_s_9590 | Ca_Nitrospira_inopinata | genome available | PMID=26610024 |
| midas_s_211 | Ca_Nitrospira_nitrosa | genome available | PMID=26610025 |
| midas_g_1393 | Ca_Obscuribacter | genus support (>94.5% id) for 3/3 (100%) MiDAS2 seqs | PMID=28365734 |
| midas_f_176 | Ca_Promineofilaceae | genome available | PMID=26905629 |
| midas_g_176 | Ca_Promineofilum | genome available | PMID=26905629 |
| midas_s_658 | Ca_Promineofilum_breve | genome available | PMID=26905629 |
| midas_s_2048 | Ca_Propionivibrio_aalborgensis | genome available | PMID=27458436 |
| midas_s_728 | Ca_Saccharimonas_aalborgensis | genome available | PMID=23707974 |
| midas_g_425 | Ca_Sarcinithrix | genus support (>94.5% id) for 19/19 (100%) MiDAS2 seqs | PMID=28365734 |
| midas_g_471 | Ca_Villigracilis | genus support (>94.5% id) for 16/16 (100%) MiDAS2 seqs | PMID=28365734 |
| midas_s_48 | Nitrospira_defluvii | genome available | PMID=20624973 |
| midas_f_57 | Rhodobacteraceae | the genus was renamed | PMID=26203335 |
| midas_g_5 | Tetrasphaera | manual curation | none |

**Table S4.** List of bacterial names with differences between genus name and type strain name inherited from Silva_132_SSURef_Nr99 (Quast et al., 2013).

| MiDAS 3 genus name | MiDAS 3 species name |
|---|---|
| Subdoligranulum | Faecalibacterium_prausnitzii |
| Enhydrobacter | Moraxella_osloensis |
| Iamia | Aquihabitans_daechungensis |
| Luteimonas | Lysobacter_lycopersici |
| Agathobacter | Eubacterium_rectale |
| Tepidimicrobium | Clostridium_ultunense |
| Eubacterium_nodatum_group | Aminicella_lysinilytica |
| Fusicatenibacter | Clostridium_clostridioforme |

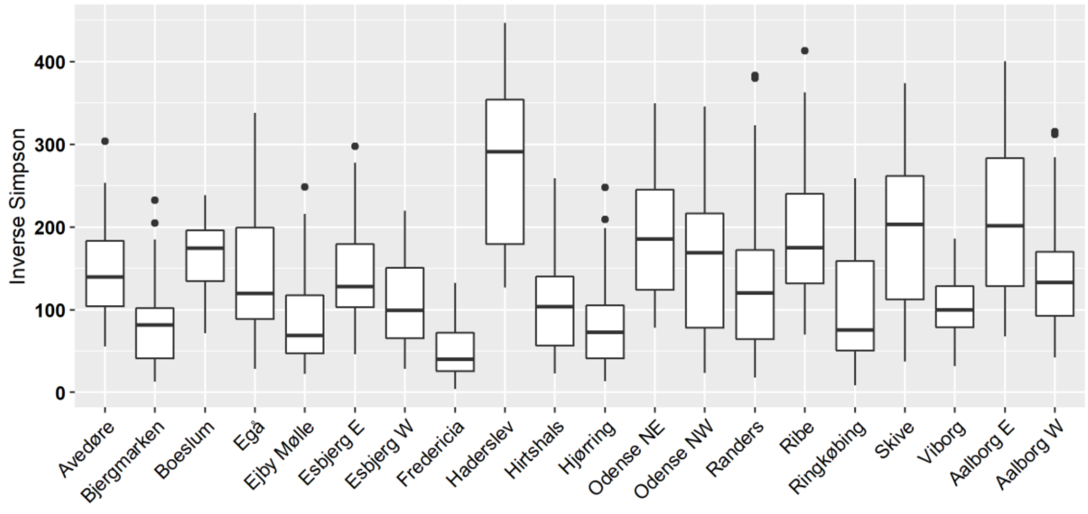


**Figure S1**. Alpha diversity estimate of activated sludge community composition using ASV-level taxa in individual plants, calculated using the Inverse Simpson index. Data represents 712 samples from 20 Danish full-scale WWTPs collected from 2006 to 2018 (with 17-51 samples per plant).

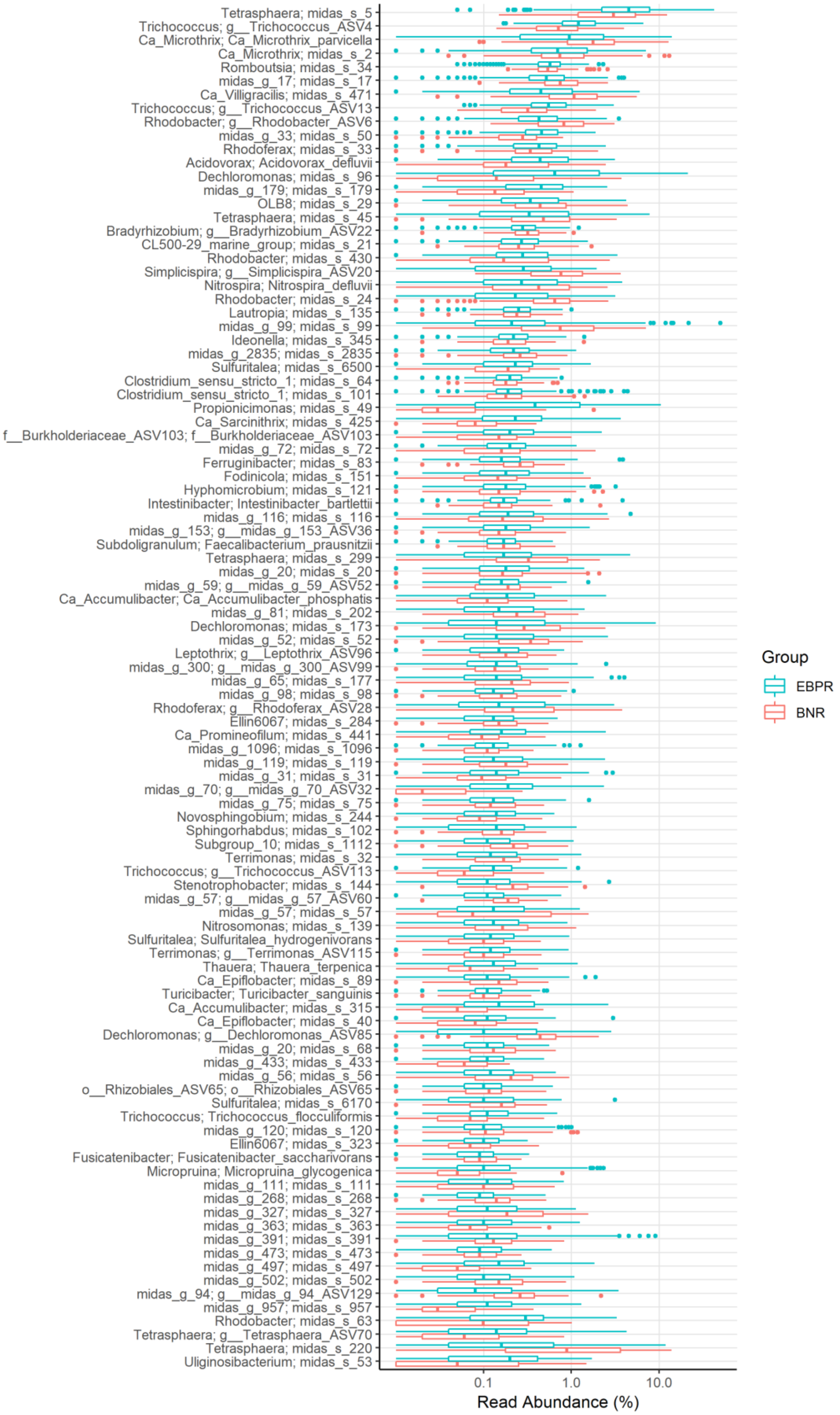


**Figure S2.** Boxplot showing the occurrence of top 100 most abundant species in Danish EBPR and BNR WWTPs.
